# Motion-in-depth perception and prey capture in the praying mantis

**DOI:** 10.1101/502583

**Authors:** Vivek Nityananda, Coline Joubier, Jerry Tan, Ghaith Tarawneh, Jenny CA Read

## Abstract

Perceiving motion-in-depth is essential to detecting approaching or receding objects, predators and prey. This can be achieved using several cues, including binocular stereoscopic cues such as changing disparity and interocular velocity differences and monocular cues such as looming. While these have been studied in detail in humans, only looming responses have been well characterized in insects and we know nothing about the role that stereo cues play and how they might interact with looming cues. We used our 3D insect cinema in a series of experiments to investigate the role of the stereo cues mentioned above, as well as looming, in the perception of motion-in-depth during predatory strikes by the praying mantis. Our results show that motion-in-depth does increase the probability of mantis strikes but only for the classic looming stimulus, an expanding luminance edge. Approach indicated by radial motion of a texture or expansion of a motion-defined edge, or by stereoscopic cues, all failed to elicit increased striking. We conclude that mantises use stereopsis to detect depth but not motion-in-depth, which is detected via looming.

**Summary Statement:** Both stereoscopic visual cues and changing luminance cues can enable detection of approaching objects. We show that mantises use looming rather than stereo cues to detect motion-in-depth.

## Introduction

Depth perception is a vital requirement for visually behaving animals. It is fundamental to be able to avoid collision with the environment or other animals. It is also important to determine how close a predator or prey is. For predatory animals, precise distance estimation is especially important in order be able to successfully execute the interception and capture of prey. Several cues could enable the perception of depth. These include cues provided by motion, such as optic flow or motion parallax, pictorial cues (such as shading and relative size) and stereoscopic cues (Nityananda and Read, 2017). The latter involve cues that convey depth as a result of comparing the differential visual input and scenes perceived by the two eyes.

A key aspect of depth perception for both predators and prey is the ability to detect motion-in-depth, i.e., when an object is approaching or receding. This would for example, be important for prey to take evasive action when predators are moving towards them. Similarly, predators would be able to use motion-in-depth to better capture prey as they come near. Just as with depth perception, several cues could contribute to the perception of motion-in-depth.

Two of the motion-in-depth cues that have received the most attention in humans are binocular: *changing disparity* (CD) and *interocular velocity differences* (IOVDs) (Cormack et al., 2017). Stereoscopic disparity refers to the difference in the position of an object as seen by the two eyes. This disparity reflects the distance to an object. Thus as an object approaches, the disparity between the two views changes. This changing disparity cue suffices to create a perception of motion-in-depth for human observers, even in the absence of other cues (Cumming and Parker, 1994). Approaching objects would also have differing velocities in each eye. For example, an object approaching along the midline would have a rightward velocity in the left eye and a leftward velocity in the right eye. These interocular velocity differences have also been shown in humans to contribute to judgements of motion-in-depth (Shioiri et al., 2000). The relative strength of the two cues depends on the precise stimulus and task; for example, IOVDs dominate for stimuli with high speeds covering wide areas of the visual field, whereas CD cues dominate for lower speeds in the central visual field (Cormack et al., 2017; Czuba et al., 2011; Parker et al., 1996)

A powerful monocular cue to motion-in-depth is looming: the increase in an object’s apparent size as it approaches. This is a special case of the more general optic flow cue to depth: when our visual scene moves directly towards us, we experience a radial flow field in which all features move radially away from the fovea. Looming cues, typically of a dark object against a light background, have been well studied in insects, where species including locusts and mantises have been shown to have escape or defensive responses to looming stimuli (Rind and Simmons, 1992; Santer et al., 2005; Yamawaki and Toh, 2009). Looming-sensitive neurons, i.e. neurons which preferentially respond to looming stimuli compared to receding or translating stimuli, have been identified in these species. In mantises, these neurons have also been implicated in defensive responses (Sato and Yamawaki, 2014; Yamawaki, 2011).

Humans use multiple cues to depth and combine them in complex ways depending on the stimulus and task (Cormack et al., 2017; Regan et al., 1979). Both monocular and binocular cues are important for humans but changing disparity often dominates perception when present (Nefs et al., 2010). Looming and changing disparity, however, both act independently upon a common stage in perception to convey a perception of motion-in-depth, so for example the two cues can cancel each other out (Regan and Beverley, 1979). In general, when multiple cues are present, individuals appear to interpret stimuli such that there is least conflict between different sources of information about motion-in-depth (Brenner et al., 1996).

Much less is known about how insects, with their far simpler nervous systems, combine multiple cues and reconcile conflicts. Praying mantises are particularly interesting animals to consider. When it comes to predation, they have an especially clear behaviour indicating their perception of depth – a predatory strike that involves a rapid extension of their forelimbs to capture prey, released only when prey is within catch range. Mantises are sensitive to multiple cues to depth including stereo cues and motion parallax due to self-motion (Nityananda et al., 2016b; Poteser and Kral, 1995; Rossel, 1983). For motion-in-depth specifically, their sensitivity to looming has been studied in some detail (Sato and Yamawaki, 2014; Yamawaki, 2011), but other cues have not been examined. In addition, looming has been studied in the context of defensive responses and has not been implicated in prey capture so far. This suggests the hypothesis that mantises use looming to avoid predators and stereopsis to catch prey. This is supported by the fact that praying mantises are the only invertebrates known to have stereoscopic vision. They use this to judge prey distance (Nityananda et al., 2016b; Rossel, 1983) and also to modulate their preference for prey size (Nityananda et al., 2016a). It is thus also possible that they exploit stereoscopic cues to motion-in-depth. Our recently-described insect 3D cinema allows us to manipulate stereo cues freely (Nityananda et al., 2016b), enabling us to investigate this question. We therefore ran a series of experiments aimed at uncovering which cues mantises use to detect the motion of prey in depth, and how these are combined.

## Results and Discussion

### Experiment 1: Briefly pulsed stereoscopic motion-in-depth cues do not influence mantis striking

We began with the random-dot stimulus exploited in our previous paper (Nityananda et al., 2018). This consists of dense, random patterns of small dark and bright dots. We have shown previously that if a patch of dots moves around such an image, mantises use the binocular disparity of the moving patch to work out whether it is in strike range (and attack if so). Furthermore, mantises continue to use binocular disparity in this way even if no dots physically move, but just invert their contrast briefly as a notional “patch” moves over them (Nityananda et al., 2018). We concluded in this earlier work that praying mantis stereopsis is fundamentally different to human stereopsis. Human stereopsis is based on the pattern of luminance (light and dark features) in the two eyes. It assumes that the two eye’s patterns are locally related by a shift and seeks to extract this disparity. Mantis stereopsis, in contrast, is completely insensitive to the detailed pattern of luminance, and is unimpaired even when the patterns in the two eyes are uncorrelated. Rather, mantis stereopsis appears to look for regions in each eye where the image is changing, and then uses the disparity between these regions.

Here, we developed a version of this stimulus which enabled us to compare constant disparity with changing disparity / interocular velocity difference. Each eye sees a different random dot pattern. A notional circular patch, corresponding to the simulated prey, spirals around the screen. As this patch passes over each dot, the dot jumps horizontally; when the patch moves off the dot, it jumps back to its old position. Thus no dots physically move around the pattern, but a ripple in the dot pattern spirals around the screen. In the Constant Depth condition, the direction of the jump was the same in both eyes (Fig. 1), so there was no interocular velocity difference. However the location of the jumping dots was offset in the two eyes, with either “crossed” disparity indicating that the patch was 2.5 cm in front of the animal, or “uncrossed” not consistent with any distance. Because the dots jumped in the same direction, the disparity of the virtual patch remains constant as it moves around the screen. In each eye at any given moment, the jumping dots define a location where the image is changing. This location moves in each eye, but the disparity between the left and right locations remains constant.

**Fig. 1.**
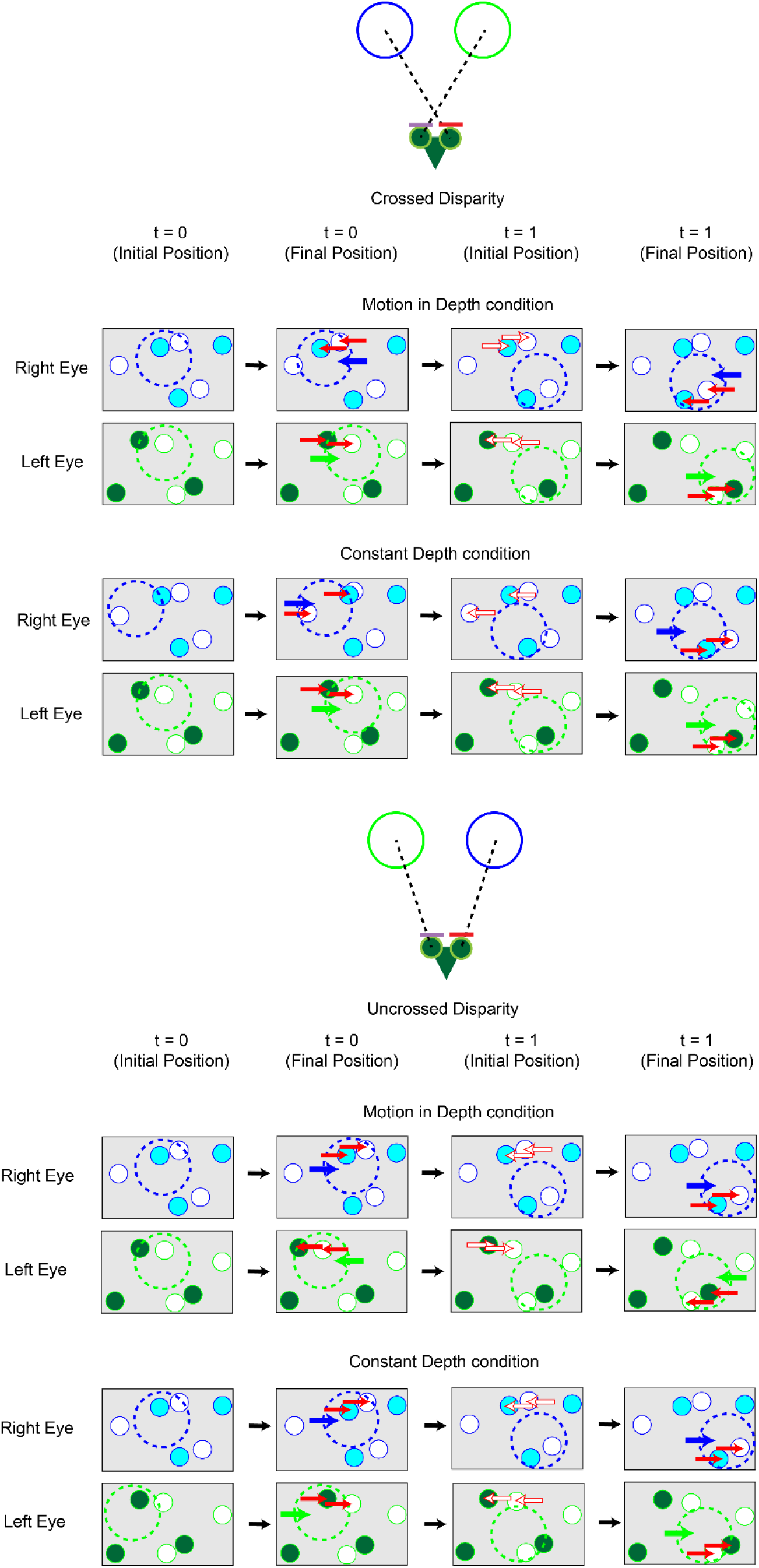
Cartoon of the stimuli in Experiment 1. The random dot stimuli consisted of uncorrelated dark and light dots against a grey equivalent background. In the Motion-in-depth (MID) condition dots jumped in opposite directions when a focal region (circles with dashed outlines) passed over them. In the Constant Depth condition, the dots jumped in the same direction. The final positions and disparities of the focal regions in both conditions were the same. Dots returned to their original position once the focal regions moved on. Both conditions were presented with crossed and uncrossed disparities. These are illustrated above - the lines of sight to the focal areas crossed in the crossed disparity conditions and didn’t cross in the uncrossed condition. In this figure, time “t” indicates a frame of the stimulus. “Final Position” indicates the visible stimulus shown on each frame; the “Initial Position” are conceptual positions shown here for clarity. At the beginning of each frame, we update the focal region location (dotted ring) according to its spiral trajectory around the screen, and any dots that are no longer in the focal region jump back to their original positions (red open arrows, Initial Position). Next, any dots that have newly entered the focal region jump horizontally, so the entire focal region effectively jumps (red arrows for individual dots, blue and green arrows for focal regions, Final Position). In the examples illustrated above, the focal regions in the Constant Depth condition move right and their final positions are matched in the MID condition. Experiments were also run where the focal regions in the Constant Depth condition moved left and the initial and final positions of the focal regions in the MID condition were accordingly shifted to match these positions. Dot size and density are chosen for clarity of illustration – see Methods for actual values.

In the Motion-In-Depth (MID) condition (Fig. 1), dots jump in opposite directions in the two eyes. Thus there is a brief pulse of interocular velocity difference as the patch moves over each region. At each moment, the location within which dots jump is identical in the two eyes, but since they jump in opposite directions, the end-point of the jump is offset in the two eyes. In the crossed disparity condition, this offset has disparity indicating 2.5 cm, so effectively there is an MID cue specifying an approach from 10 cm (the screen plane) to 2.5 cm. Conversely in the uncrossed disparity condition, the binocular cues imply a recession. Critically, the monocular stimuli are individually indistinguishable in the Constant Depth and MID conditions.

The results are shown in Fig. 2. Consistently with our previous work, mantises robustly discriminated depth in the Constant Depth condition. All six mantises struck more for crossed disparity and this difference was significant at the population level (stats below). In contrast, for the MID condition, three of six mantises did not strike at all. Of the three that struck, two struck more for the crossed condition and one for uncrossed, so that overall there is no difference between crossed and uncrossed.

**Fig. 2.**
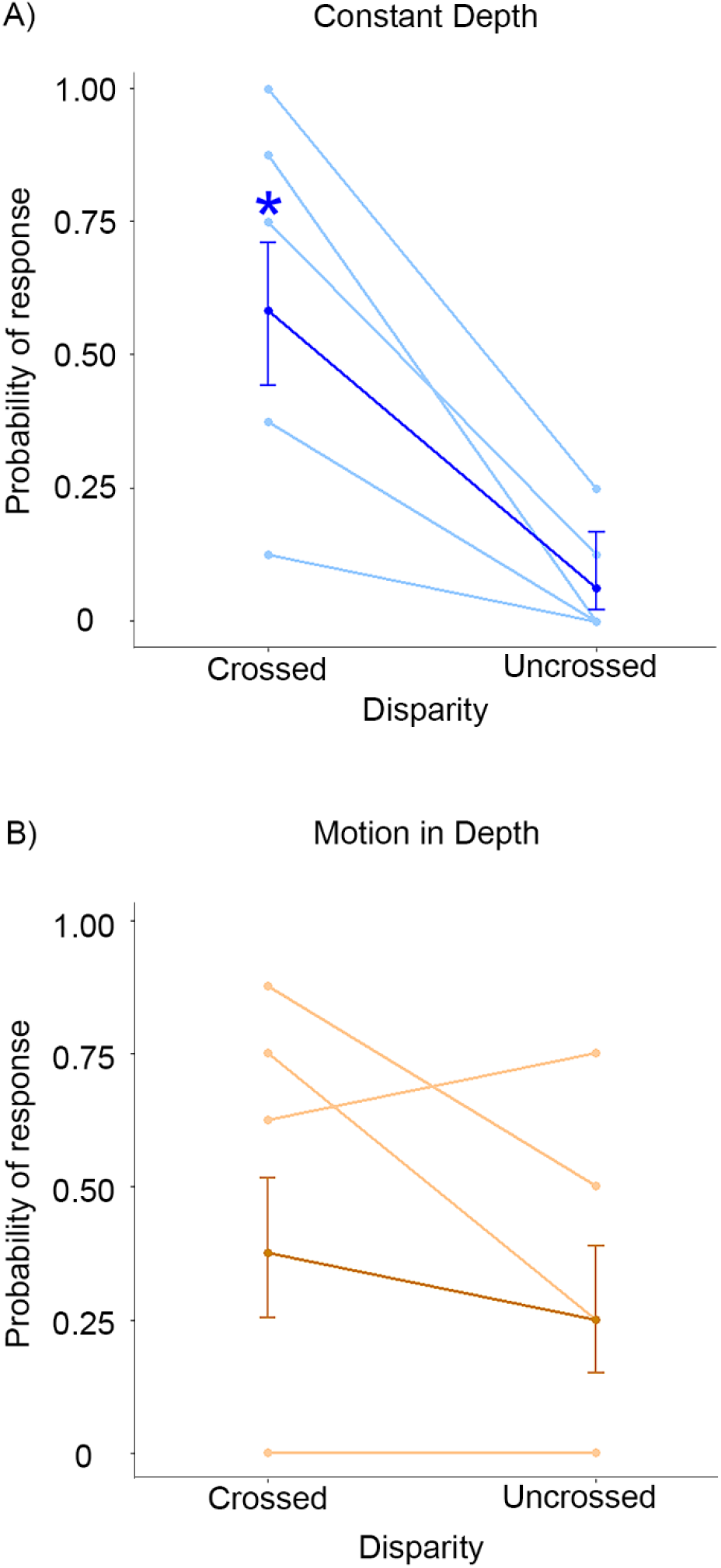
Probability of mantis strikes in response to random-dot stimuli stimuli a) with constant depth and b) with motion-in-depth. Stimuli consisted of a random-dot background with a targets defined by a spiralling focal area defined by dots in each eye briefly jumping in either A) the same direction (i.e. without IOVD and changing disparity cues) or B) opposite directions (i.e. with IOVD and changing disparity cues). The disparity of targets was either crossed, and indicated a target 2.5 cm from the mantis at the final position or uncrossed, where the final position of the targets had the same parallax as the crossed condition but with the left and right positions swapped on the computer screen. Bold lines indicate the mean strike probabilities across all six animals with 95% binomial confidence error bars. Lighter lines represent the response of individual animals. Asterisks mark statistically significant increases (P < 0.05).

The model that best explained our results included an interaction between the MID and Disparity factors (Interaction Model BIC = 173.4; Non-interaction Model = 184.5). Both the MID condition and Disparity had a significant main effect on the probability of strikes (MID: Estimate =2.0198, P = 0.007641; Disparity: Estimate = 4.6592, P=7.87e^−8^). Mantises were likely to strike at crossed disparities (crossed disparity mean strike probability= 0.4791667; uncrossed disparity mean strike probability = 0.15625) and in the Constant Depth condition (Constant Depth mean strike probability= 0.3229167; MID mean strike probability = 0.3125) (Fig. 2). There was also a significant interaction between the two factors (Estimate = −3.6934, P = 0.000219). This interaction shows that mantises were significantly less likely to strike when the stimulus was presented with Constant uncrossed disparity (Mean probability = 0.0625) compared to Constant crossed (Mean probability = 0.5833) but in the MID condition, they struck equally for change in either direction (Crossed mean probability = 0.375, and Uncrossed mean probability = 0.25).

Clearly, the stereoscopic cues to motion-in-depth in this stimulus were either not detected by the mantis visual system, or did not influence the decision to strike. Thus Experiment 1 provides no evidence that praying mantises can exploit binocular cues to motion-in-depth.

### Experiment 2: Persistent, veridical stereoscopic motion-in-depth cues also do not influence mantis striking, but looming does

In Experiment 1, the stereoscopic motion-in-depth cues were presented only very briefly, and were not consistent with the approach of a real object. It would therefore be premature to conclude from Experiment 1 that the mantis visual system cannot exploit stereoscopic motion-in-depth cues in more naturalistic stimuli. To this end, we returned to a more naturalistic stimulus which we have previously found readily elicits strikes (Nityananda et al., 2016a; Nityananda et al., 2016b). This consists of a dark disk spiralling round on a brighter background. In our previous experiments, the disk had a constant screen parallax, designed to depict an object at 2.5 cm when presented with “crossed” geometry, and constant size. We now explored changing the parallax and screen size during the stimulus presentation, so as to depict an object approaching at a constant speed (condition 1 in Fig. 3). We found that a certain amount of spiral motion was still necessary in order to elicit enough strikes for analysis; the mantises did not respond to an object approaching head-on. Thus Condition 1 depicted an object spiralling in the frontoparallel plane (X, Y) while approaching from Z=20 cm to Z=2.5 cm. In our other conditions, we held either the size or disparity fixed at a single value (Fig. 3).

**Fig. 3.**
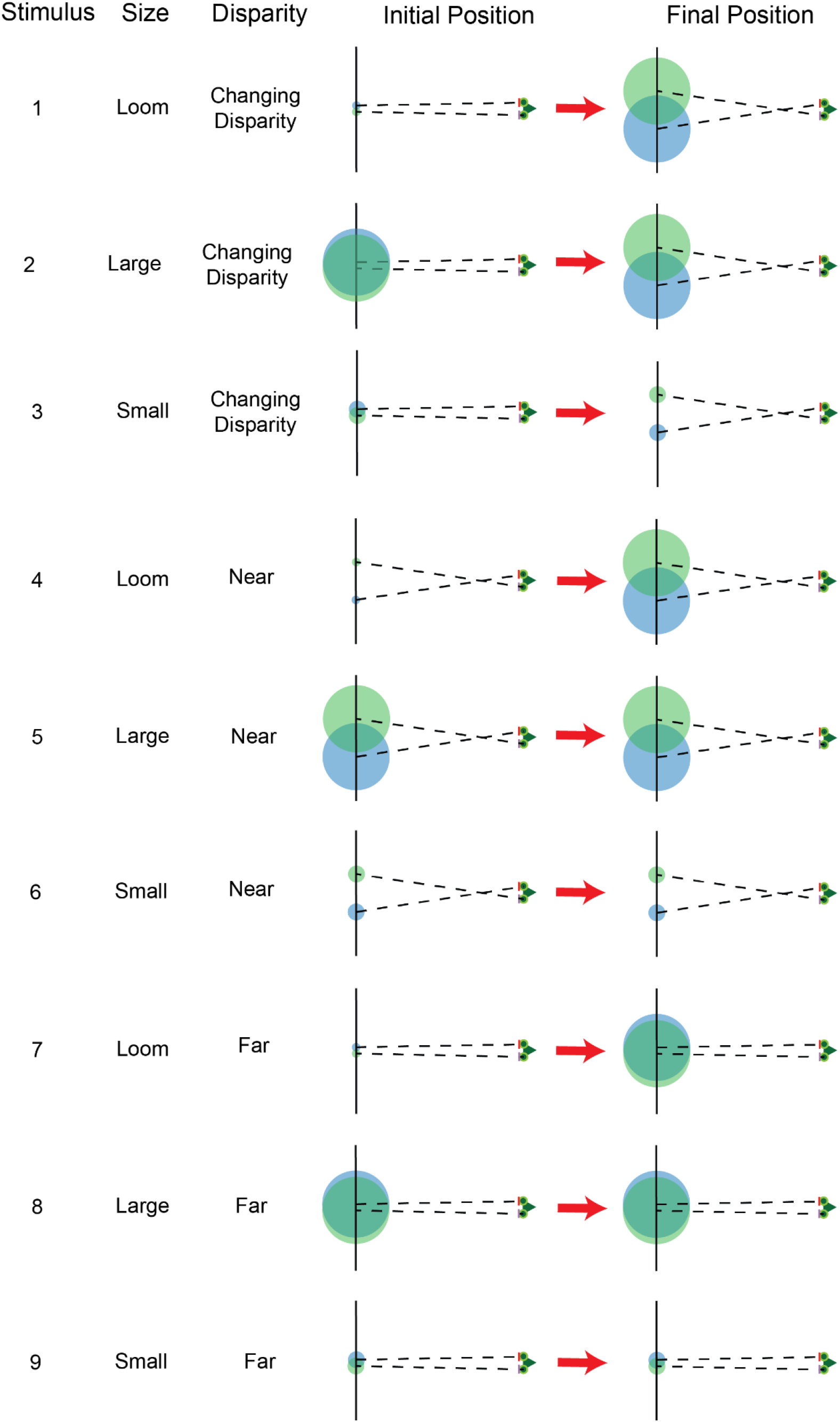
Stimuli presented in Experiment 2. The stimulus here was a dark spiralling disc presented against a grey background. The disparity between the discs seen in each eye and the sizes of the discs were varied in different stimuli to present different combinations of size and disparity. Mantis head with 3D glasses depicted on the right with dotted lines of sight reaching to the centre of the discs presented on the screen which was 10 cm away. Initial and final states of each stimulus demonstrate changes in size or disparity or lack thereof.

Results are shown in Fig. 4. Our veridical stimulus (condition 1 in Fig. 3) contained both looming and stereoscopic motion-in-depth cues. It increased in size, its disparity changed, and as a consequence the interocular velocity differences also changed consistent with its approach. This stimulus elicited strikes on around 60% of trials on average. We then explored removing either looming or stereoscopic cues to motion-in-depth. Conditions 2 and 3 remove the looming cue; now the angular size remained constant (either large, consistent with a nearby object, or small, consistent with a distant one) although the stereoscopic cues still specified an approaching object. Both these elicited fewer strikes, although the large fixed-size object was clearly preferred to the small object. The remaining 6 conditions investigate the response when stereoscopic cues specify a constant distance. In the “near disparity” stimuli (4–6), the target spirals at a constant stereo-defined distance of 2.5 cm from the mantis. The responses were similar to those to the “changing disparity” stimuli (1–3): once again strikes are elicited most when the looming cue is present, less when the angular size is constant and large, and least of all when it is constant and small. In the “far disparity” stimuli (7–9), where stereo cues indicate that the prey is at a constant distance of 10 cm, out of strike range, the relative proportions are similar but the overall strike rate is – not surprisingly – greatly reduced.

**Fig. 4.**
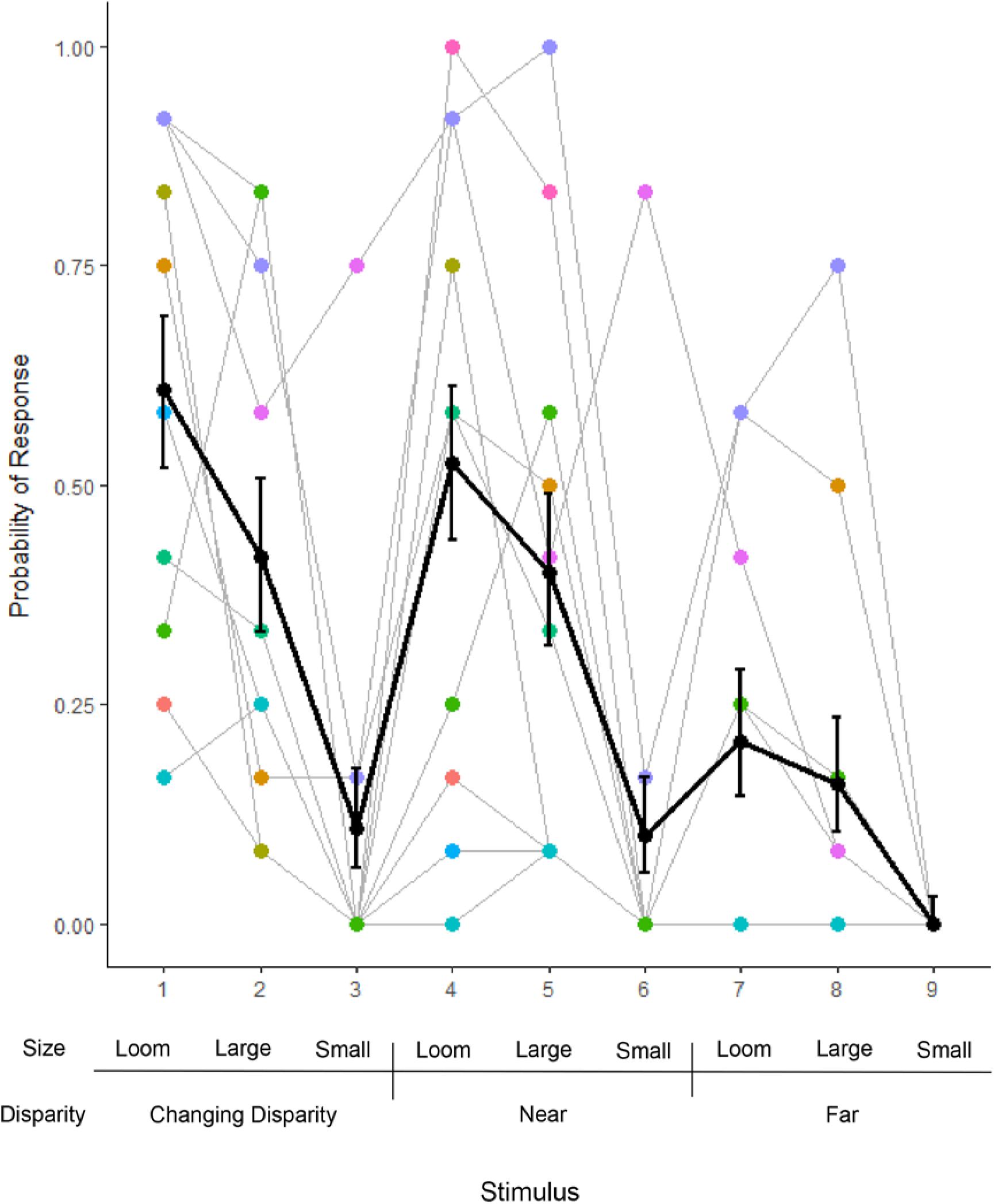
Mantis probability of striking to different conditions with or without motion-in-depth. Bold black line indicates the mean probability across all ten animals with 95% binomial confidence error bars. Lighter lines represent the response of individual animals with animal identity indicate by the colour of the points. The different size and disparity combinations for each stimulus condition are indicated below. ‘Near’ disparities simulated a target 2.5 cm from the mantis and ‘Far’ disparities simulated a target 10 cm from the mantis. ‘Large’ and ‘Small’ sizes subtended the same visual angle as a target 2.5 cm (22.62°) and 10 cm (5.72°) respectively from the mantis. ‘Loom’ and ‘Changing Disparity’ conditions simulated a target approaching the mantis from a distance of 20 cm to a distance of 2.5 cm with size and disparity cues respectively. In this experiment, the target was a luminance-defined dark target against a light background.

The model that best explained our results included both Size and Disparity as factors without interaction effects (without interaction: BIC = 883.3; with interaction: BIC = 905.2). In this model three of the levels had significant effects. These were Size: Small (Estimate = −2.5004, P < 2e^−16^), Size: Loom (Estimate = 0.7993, P = 3.76e^−5^) and Disparity: Far (Estimate = −1.9733, P = 1.35e^−15^) (Fig. 4). This shows that mantises are less likely to strike at a target whose disparity indicates it is 10 cm away (out of catch range) or if it subtends a smaller angle of 5.72°. Looming, however, significantly increases the chances of a strike. If angular size is constant, mantises prefer our large prey (22.62°) to our small prey (5.72°). However, they have an even greater preference for prey whose angular size changes from small to large. Since such angular changes in the real world are almost always caused by approach, this implies that mantises preferentially attack approaching objects. This is the first evidence that mantises use looming information when hunting their prey, and not only to detect the approach of a predator (Sato and Yamawaki, 2014; Yamawaki, 2011).

Importantly, the changing-disparity cue did not have a significant effect on the probability of striking (Estimate = 0.2590, P = 0.194), although far disparity significantly suppressed striking (Estimate = −1.9733, P = 1.35e^−15^). That is, mantises are more likely to strike when stereopsis indicates that an object is in catch range, but this preference is not stronger when stereopsis indicates that the object is approaching. This implies that, although mantises preferentially attack approaching objects, and although they possess stereoscopic information about object distance, they do *not* use stereoscopic information to detect *changes* in distance. If they did, their preference for approaching objects would mean that they would be even more likely to strike when disparity indicated an approaching object was now within catch range than if an object simply moved at a constant distance within catch range.

Thus, Experiment 2 implies that mantises use monocular looming cues to detect approaching objects, and stereoscopic disparity to tell whether an object is in catch range. However, it implies that mantises do not use stereoscopic motion-in-depth cues, whether changes in disparity or interocular velocity differences, to detect approaching objects. This is consistent with our conclusions from Experiment 1.

### Experiment 3: Looming cues require a luminance edge

Experiments 1 and 2 both imply that mantises use stereopsis to detect depth, but not motion-in-depth. Experiment 2 confirms previous literature that they do use looming to detect motion-in-depth. As noted in the Introduction, looming is a special case of optic flow cues to motion relative to the environment. When one moves towards an object or surface, or it moves towards you, points on the surface flow radially across the retina. The term “looming” is generally reserved for a dark object increasing in size, as in our Experiment 2. This produces a radial expansion of a high-contrast luminance edge, without any radial motion beyond the edge. Here, we wanted to ask if this moving luminance edge is required for motion-in-depth perception in mantis predation. We envisaged various possibilities, namely the mantis visual system detects the approach of a prey item if:

i. There is expanding radial first-order motion of a luminance boundary.
ii. There is expanding radial first-order motion, but not of a luminance boundary.
iii. There is expanding radial motion of a second-order boundary, but without first-order motion.

As we have seen, an example of case (i) is the expanding dark disk, which we showed in Experiment 2 does contribute to motion-in-depth perception in mantis predation. An example of case (ii) is an expanding star field, as when the USS Enterprise enters warp. This is a familiar stimulus in the optic flow literature, but to our knowledge has not been investigated in predation. For an approaching prey object, the radial expansion would be confined to a small part of the visual field, corresponding to the prey. This sort of stimulus could occur if the prey had the same mean luminance as the background, but had patterning on its body which would produce radial flow when the prey moved towards the mantis. Case (iii) is motivated by our finding that mantis stereopsis does not require first-order motion (as in the “luminance-flip” stimulus in (Nityananda et al., 2018)). This made us wonder if mantises might also be sensitive to the expansion of a boundary without any first-order motion.

To test the latter two cases, we used a random-dot pattern like that in Experiment 1. As before, a notional patch spiralled around the screen. To provide expanding radial first-order motion without an expanding luminance boundary (case (ii)), when the patch passed over a dot, the dot began to move radially away from the centre of the patch (Fig. 5, top row). When the dot passed over the edge of the patch, it vanished. This stimulus thus contained radial motion within the patch, similar to that which would be provided if the patch was approaching, and a motion-defined boundary defined by where the moving dots vanished. We further distinguished “looming” and “non-looming” versions of this stimulus. In the non-looming version, the patch stayed the same size as it spiralled around the screen, and disparity remained constant at a value implying an object in catch range, i.e., 2.5 cm from the screen. In the “looming” version, the patch increased in size (i.e. the motion-defined boundary expanded radially), and the patch’s disparity also changed. Thus in the “looming” version, monocular motion-in-depth cues are potentially available both from the radial motion of dots themselves and also from the radial motion of the motion-defined boundary as well as from the stereoscopic motion-in-depth cues; in the “non-looming” version, the only motion-in-depth cue is the radial motion of the dots.

**Fig. 5.**
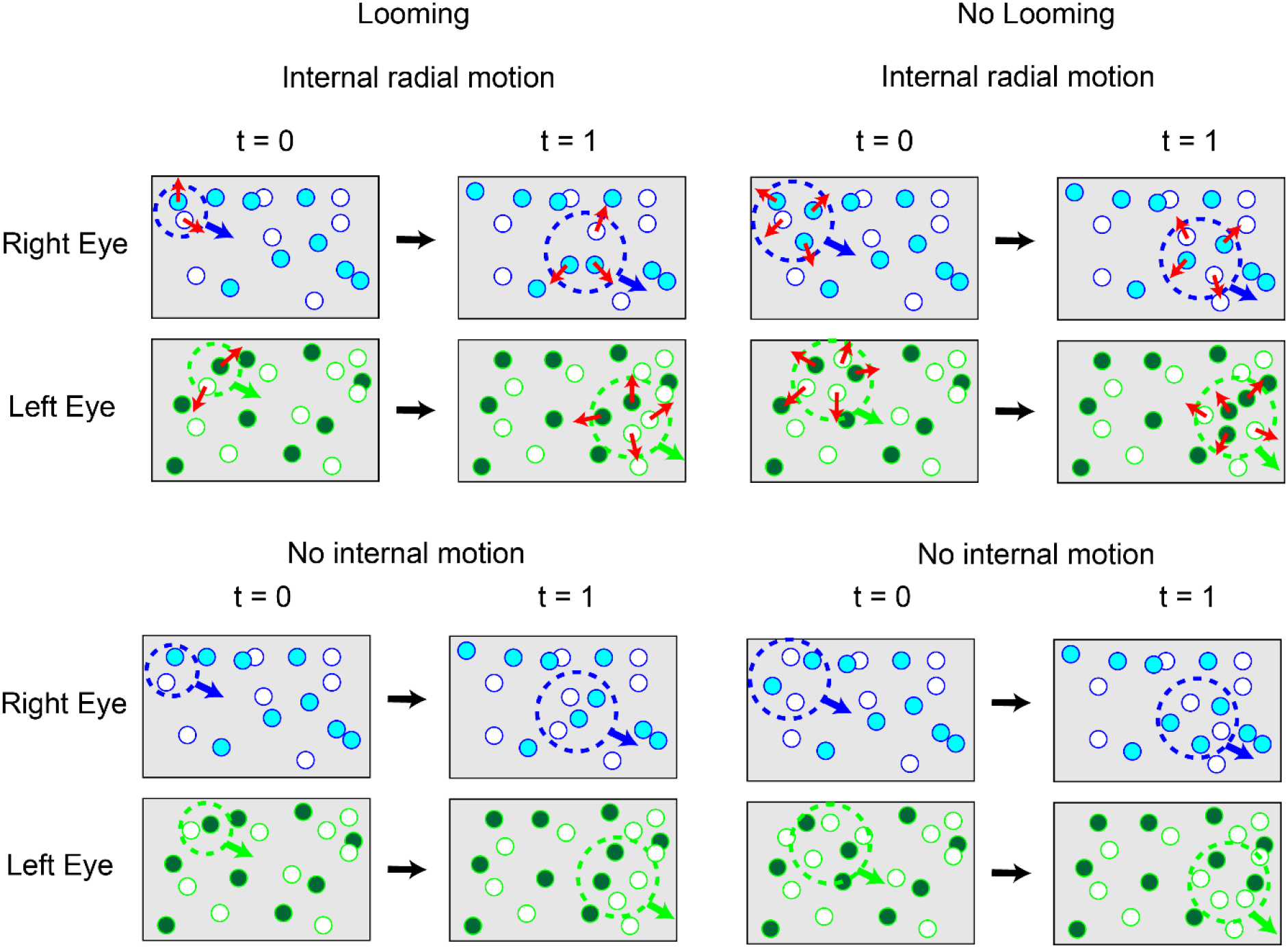
Cartoon of the stimuli in Experiment 3. The random dot stimuli consisted of uncorrelated dark and light dots against a grey equivalent background. The target spiralled in (dotted circles and blue and green arrows) over the dots with either expanding radial motion of the dots within (red arrows) or no motion of the dots. In the looming condition, the target expanded over the course of the stimulus presentation while in the non-looming condition, the target stayed the same size as the final size in the looming condition. All stimuli were presented with both crossed and uncrossed disparity but only the crossed disparity case is depicted here. Dot size and density are chosen for clarity of illustration – see Methods for actual values.

For case (iii), we used a second-order motion stimulus (Fig. 5, bottom row). Now, when the patch entered a region of the screen, the dots in that region vanished and were replaced with a different random dot pattern. When the patch moved away, the new dots vanished and the original dots returned. This type of stimulus is called “drift balanced” (Chubb and Sperling, 1988). The appearance and disappearance of dots at the boundary of the patch provides a second-order motion cue to the motion of the patch. Again, we tested “looming” and “non-looming” versions of this stimulus. In the non-looming version, the patch stayed the same size and the disparity stayed constant. There were thus no motion-in-depth cues at all. In the looming version, the patch increased in size and the disparity changed. There was thus a monocular motion-in-depth from the radial motion of the second-order boundary, as well as the stereoscopic cues.

This complicated set of conditions is summarised in Table 1 and the results are shown in Fig. 6. The model that best explained our results did not have an interaction effect between the factors (Without Interaction BIC = 827.2; With Interaction BIC = 845.7). Disparity had a significant effect on the probability of striking (Estimate = 2.6939, = 0.005511), with crossed disparity stimuli resulting in more strikes than uncrossed stimuli. Looming had a significantly negative effect on the probability of striking (Estimate = −0.6761, P = 0.000219). Contrary to the results in Experiment 1, a looming stimulus defined by a motion-edge thus reduced the probability of striking compared to a non-looming stimulus (Looming mean strike probability= 0.4464286; Non-looming mean strike probability = 0.5379464). Motion Condition, did not have a significant effect on the probability of striking (Estimate = −0.3473, P = 0.054882) (Fig. 6). There was thus no difference if the motion-edge was defined by internal outward motion (Fig. 6A) or a motion boundary without internal motion (Fig. 6B).

**Table 1.** Cues provided by the various stimulus conditions in Experiment 3. Asterisks and red text indicate conditions that produced a significant increase in strike rate.

| Condition |  |  | Motion-in-depth cues |  |  |
| --- | --- | --- | --- | --- | --- |
|  |  |  | Monocular |  | Stereoscopic |
| Motion condition | Looming condition | Disparity condition | Radial motion of dots | Radial motion of motion-defined boundary | Changing disparity and interocular velocity difference |
| First-order motion | Looming | Crossed* | Outward, indicating approach | Outward, indicating approach | Both indicate approach |
|  |  | Uncrossed | Outward, indicating approach | Outward, indicating approach | Both indicate recession |
|  | Non-looming* | Crossed* | Outward, indicating approach | Fixed, indicating constant distance | Both indicate constant distance (disparity indicates 2.5 cm) |
|  |  | Uncrossed | Outward, indicating approach | Fixed, indicating constant distance | Both indicate constant distance (disparity is uncrossed) |
| Second-order motion | Looming | Crossed* | None, indicating constant distance | Outward, indicating approach | Both indicate approach |
|  |  | Uncrossed | None, indicating *constant distance* | Outward, indicating approach | Both indicate recession |
|  | Non-looming* | Crossed* | None, indicating *constant distance* | Fixed, indicating *constant distance* | Both indicate *constant distance* (disparity indicates 2.5 cm) |
|  |  | Uncrossed | None, indicating *constant distance* | Fixed, indicating *constant distance* | Both indicate *constant distance* (disparity is uncrossed) |

**Fig. 6.**
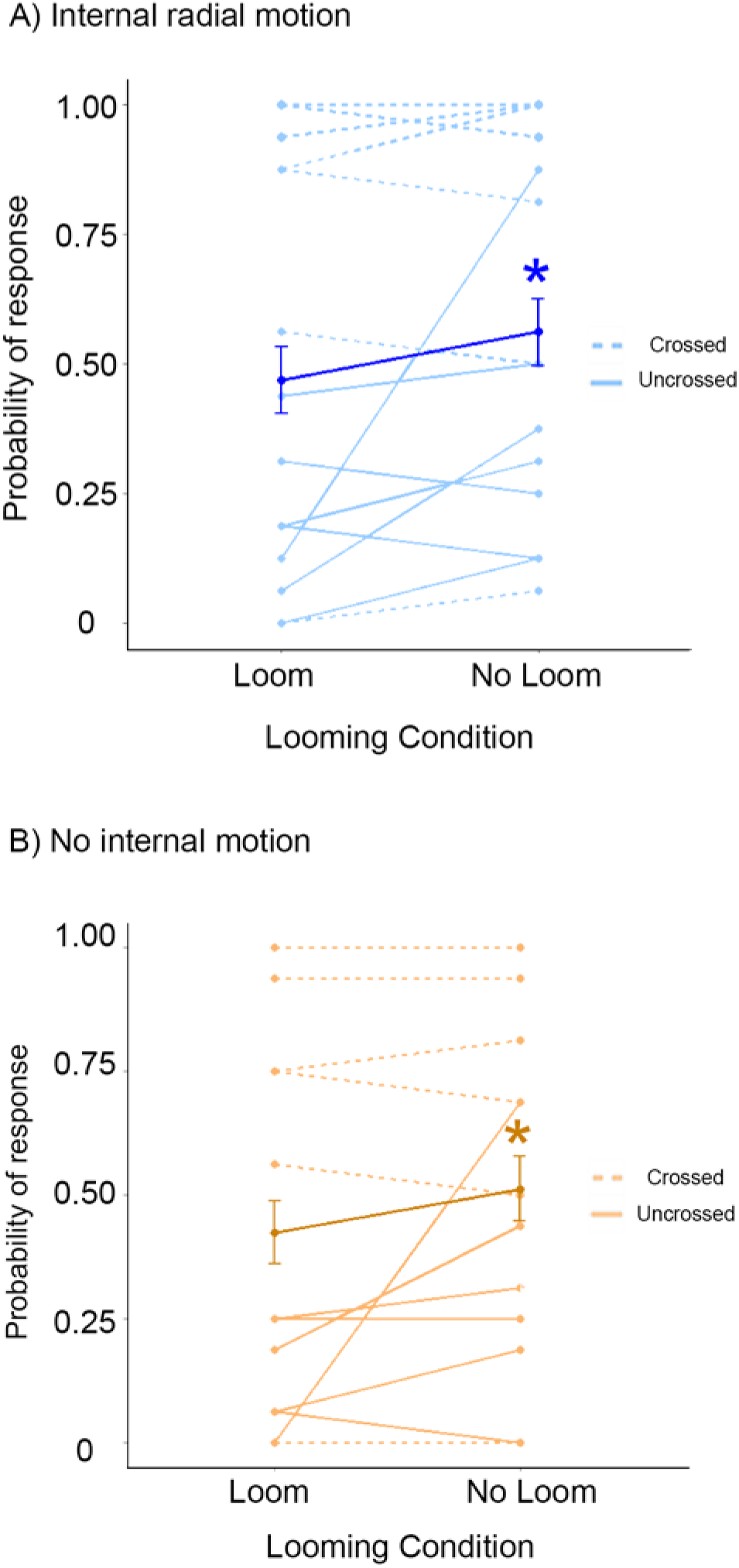
The probability of mantises striking in response to targets with motion-defined looming. Stimuli consisted of a random-dot background with a target defined by a spiralling focal area defined by two different forms of motion: A) dots streaming outwards from the centre of the target and B) a motion boundary defined by the target being a moving window to a different pattern of dots (drift-balanced motion). Both motion conditions were presented with the target either looming or with a fixed size. In the looming condition, the target changed in size and disparity to simulate a target approaching the mantis from a distance of 20 cm to a distance of 2.5 cm in the crossed disparity stimuli. In the non-looming condition the target had a fixed size and disparity simulating a target at a distance of 2.5 cm in the crossed disparity stimuli. The uncrossed disparity stimuli in both motion conditions had the same parallax as the crossed disparity condition but with the left and right eyes swapped. Bold lines indicate the mean probabilities across all seven animals with 95% binomial confidence error bars. Different animals were used in the crossed and uncrossed conditions. Lighter lines represent the response of individual animals with dotted lighter lines representing responses in the crossed disparity condition and solid lighter lines representing responses in the uncrossed disparity condition. Asterisks mark statistically significant increases (P < 0.05).

The simplest explanation of this pattern of results is that none of these stimuli produces a percept of prey motion-in-depth. Mantises struck more for the stimuli with crossed disparity, since here the stereoscopic depth cues indicated a prey item in catch range for at least some of the trial duration (with uncrossed stimuli, the disparity indicated unattractive or undefined distances). Whereas in Experiment 2 we found that a looming cue provided by a radially expanding luminance-defined boundary produced additional increases in strike rate even for crossed disparity, here we find that no such increase is provided by a radially expanding motion-defined boundary, or by radial dot motion. The results are consistent with our previous finding that mantises do not use stereoscopic motion-in-depth cues. In this experiment, the looming stimuli increased in angular size from 5.72° to 22.62°, while the non-looming stimuli were fixed at the larger size (22.62°). The greater strike rate for non-looming stimuli thus presumably reflects mantises’ preference for larger prey (Nityananda et al., 2016a; Prete and Mahaffey, 1993), visible in Fig. 4 (greater response for “large (22.62°)” vs “small (5.72°)” non-looming stimuli).

## General Discussion

Detecting motion-in-depth is important for many purposes: for locomotion, for avoiding predation and for predation. Here, we have investigated the cues used by an insect predator, the praying mantis, to detect prey motion-in-depth. We show that mantises do detect prey motion-in-depth using looming cues and preferentially attack targets which are approaching, presumably because these are more likely to be successfully captured.

Several previous studies have shown looming cues to be important for insects, including mantises (Rind and Simmons, 1992; Santer et al., 2005; Sato and Yamawaki, 2014; Simmons and Rind, 1992; Yamawaki, 2011; Yamawaki and Toh, 2003). Detecting looming in locusts and mantises relies on certain critical cues. These include fast moving dark edges, especially when moving apart to indicate an expanding shape (Simmons and Rind, 1992; Yamawaki and Toh, 2009). Our experiment 2 reconfirms the importance of a clear luminance-defined moving edge for the perception of looming, and provides new evidence about the importance of this cue in prey detection. Previous studies of looming in mantises used a stimulus that expanded radially without any lateral motion, simulating an object directly approaching the mantis. These typically elicit a defensive response, with very few strikes (Sato and Yamawaki, 2014; Yamawaki, 2011). This is presumably because the most likely interpretation of such a stimulus is an approaching predator, intent on consuming the mantis. Our stimuli were designed to differ from the stimuli used in these past experiments to specifically ask if looming can play a role in eliciting mantis predatory strikes. Rather than using a disc that expands without any lateral motion, our stimuli follow a spiral motion path which we have previously found is particularly effective in eliciting strikes. In these stimuli, looming produced an increase in strikes. Thus, these stimuli, combining lateral motion and looming, have enabled us to show for the first time that luminance-defined looming cues to motion-in-depth are used to guide predatory behaviour in the praying mantis.

In principle, there are several other cues to motion-in-depth, including optic flow within a non-luminance-defined target, radial expansion defined by second-order motion, and the binocular cues of changing disparity and interocular velocity differences. None of these have been previously investigated in the context of praying mantis predation. The binocular cues are particularly interesting, given that the praying mantis has a wide binocular overlap and is the only invertebrate known to use stereoscopic disparity for depth perception. Thus it is fascinating to ask whether mantises can exploit their stereoscopic vision to obtain additional information about motion-in-depth.

None of our experiments provided evidence that praying mantises exploit any of these other cues to motion-in-depth. Disparity cues are certainly important in the perception of distance itself, but appear not to contribute to the perception of *changes* in distance. Indeed, we found no evidence that the mantis visual system exploits binocular cues to motion-in-depth, whether these are presented briefly and in the absence of other cues (as in Experiment 1) or over several seconds in naturalistic stimuli (as in Experiment 2). Predatory strikes are likely when stereopsis indicates the prey is within catch range (whether or not it reached there by approaching from beyond catch range), and when luminance-defined looming cues indicate that the prey is approaching. Of course, it is impossible to prove a negative, so it remains possible that mantises do exploit other cues to motion-in-depth in stimulus configurations which we have not investigated. However, for the moment, Occam’s razor suggests the conclusion that luminance-defined looming is the sole motion-in-depth cue used in praying mantis predatory behaviour. It is important to note that in nature, all three of the cues tested in this paper would usually co-occur and so in principle tracking any one of them would be sufficient to detect motion-in-depth in the overwhelming majority of natural cases. This is presumably why the mantis has not experienced selection pressure sufficient to evolve mechanisms to detect all the possible cues to motion-in-depth.

It therefore appears that mantises have two specialized modules for different functions. While the stereo system uses disparity cues to detect the depth to prey objects in a single primary plane of interest, the looming detection system is used to detect approaching objects. Both systems contribute to prey detection but looming may be particularly important for predator detection. Indeed, larger looming stimuli trigger a defensive response where the mantis withdraws its legs and freezes as one would expect in response to an approaching predator (Sato and Yamawaki, 2014). Having both systems could, for example, enable mantises to detect prey while simultaneously looking out for approaching predators. In addition, relying on monocular rather than binocular cues for motion-in-depth would allow individuals with damaged eyes or obscured fields of view to still detect and evade approaching predators.

In humans, the presence of multiple processing pathways for motion-in-depth has been argued to enable complementarity and redundancy. It could also allow for different specialized pathways for particular aspects of stimuli. It has, for example, been argued that changes in disparity help in the detection of stereo motion, while interocular velocity differences help in computing the speed of stereo motion (Brooks, 2002). Given that insect brains are orders of magnitude smaller than human brains - with less than a million neurons-it seems that they do not exploit these multiple cues to motion in depth. Rather, despite having binocular stereopsis, they rely solely on looming to detect objects’ approach.

## Methods

### Mantises

All experiments were carried out on adult female mantises of the species *Sphodromantis lineola*. Mantises were housed in semi-transparent cages measuring 7 cm by 7 cm by 9 cm and were provided a small twig to perch on. Room temperature was maintained at 25 °C. Mantises were fed one cricket three times a week, and their cages were misted at the same time. On experiment days, mantises were not fed to ensure motivation. All applicable international, national, and/or institutional guidelines for the care and use of animals were followed. All procedures performed in studies involving animals were in accordance with the ethical standards of the institution or practice at which the studies were conducted.

### 3D Glasses

All mantises were fitted with 3D glasses prior to experiments. These consisted of coloured filters cut into teardrop shapes with a maximum length of 7 cm. The filters used were LEE^®^ colour filters (http://www.leefilters.com/) with a filter of a different colour used for each eye. The LEE filters used were 797 Purple and 135 Deep Golden Amber (See Fig. S1 for spectral transmission details). Mantises were placed in their cages in a table top freezer (Argos Value Range DD1–05 Table top Freezer) for 5–8 minutes and subsequently held down using modelling clay. The glasses were then affixed onto the mantis’s pronotum using a mixture of beeswax and Rosin and a wax melter (Denta Star S ST08). A small electronic component was also fixed on the back of the mantis. This fit into a corresponding component on the experimental stand during experiments. After the glasses were fixed, the mantises were placed back in their cages and allowed to recover overnight prior to any experiments.

### Visual Stimulation

All stimuli were presented on a Dell U2413 LED monitor with custom written programs in Matlab using the Psychophysics Toolbox (Brainard, 1997; Kleiner et al., 2007). The monitor had dimensions of 51.8 × 32.4 cm (1920 × 1200 pixels) and a screen resolution of 37 pixels/cm. The refresh rate of the monitor was 60 Hz. Mantises were fixed onto a stand with the help of the component fixed onto their backs which fit into a corresponding component on the stand. Mantises were upside-down and held onto a cardboard disc with their legs. They were thus freely able to move their heads and forelegs, but the viewing distance was fixed. All stimuli were presented on a screen placed at a distance of 10 cm in front of the mantis. Stimuli were output in the green and blue channels with output in these channels weighted to adjust for the transmission of the filters (Fig. S1) and the spectral sensitivity of mantises (Rossel, 1979; Sontag, 1971). The blue visual output was 13% that of the green visual output. This ensured approximately equal input of blue and green light to the left and right visual systems after filtering through the glasses and the spectral sensitivity of the mantis. Before each experiment trial, mantises were shown a stimulus to check their motivation. This consisted of a dark disc against a grey background that swirled in from the periphery to the centre as described in Eqns. 1–5 below. This stimulus is known to reliably elicit strikes (Nityananda et al., 2016a; Nityananda et al., 2016b; Nityananda et al., 2018). Experiments were only carried out if the mantis struck at this stimulus twice in a row. Mantises were also checked for motivation in this way after experimental runs and the data were analysed only if the mantises struck at this stimulus twice in a row after the experimental run. 21 out of 91 experimental runs were excluded based on this criterion.

### Experiment 1: Briefly-pulsed Stereoscopic Depth vs Motion-in-Depth cues

In our first experiment, we focussed on stereo cues and asked whether brief interocular velocity differences (IOVDs) and changing disparity (CD) cues could contribute to motion-in-depth perception and strikes compared to motion cues in a single disparity plane with no IOVD cues. To enable us to dissect out the two conditions, we made use of random-dot stimuli. The stimulus here consisted of a grey-equivalent background with light and dark random dots that were uncorrelated between the two eyes. Each dot had a diameter of 25 pixels (subtending an average angle of 1.8° across all screen positions and an angle of 3.9° directly in front of the mantis). A focal region moved over this background with a spiral motion from the periphery of the screen to the centre as described previously (Nityananda et al., 2018). The equations describing the motion are

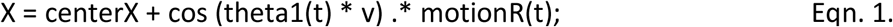

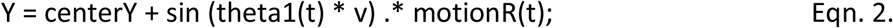

Where X and Y define the x and y positions of the centre of the target, centerX and centerY correspond to the initial x and y position of the target centre, v = 0.5 and t is the instantaneous time.

The other parameters were defined by the following equations:

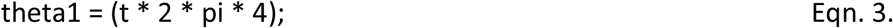

and

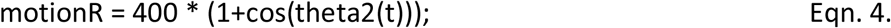

where

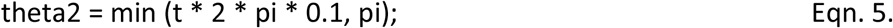

In two different conditions, dots within the focal region moved to generate IOVDs or not (Fig. 1).

In the motion-in-depth (MID) condition (Fig. 1A, C), the focal region had the same location in both eyes, i.e. zero disparity, as it swirled around the frame. When dots came within the focal region during its motion, they made a short jump in opposite directions in each eye. This jump introduced a disparity between the focal regions in each eye, equal to twice the jump size. The experiment was run in two disparity conditions – crossed and uncrossed. In the crossed disparity condition, the final position of the regions was such that the lines of sight from the two eyes to the regions visible to them crossed and the screen parallax between the regions conveyed a disparity simulating a target 2.5 cm in front of the mantis. In the uncrossed disparity condition, the final positions of the regions was the same as in the crossed condition but with left and right eyes swapped. The lines of sight to the final position of the regions did not cross and thus did not correspond to a coherent target. Thus, over the course of the motion, both IOVD and disparity cues conveyed motion-in-depth. In the crossed condition, both of them conveyed a target approaching the mantis from 10 cm away to 2.5 cm away. In the uncrossed condition, they corresponded to a target receding from 10 cm away towards infinity.

In the Constant Depth condition (Fig. 1B, D), the focal regions in each eye were separated from the start with the same screen parallax as in the final positions in the MID condition. As the regions passed over the background dots, these dots jumped in the same direction in each eye, thereby preserving the parallax and the disparity difference conveyed. Crucially, over the course of the motion, both IOVD and disparity cues thus conveyed a constant depth plane. This stimulus was also presented with both crossed and uncrossed disparity conditions. In the crossed disparity condition, the target was simulated to move laterally in a single depth plane which was 2.5 cm away from the mantis. In the uncrossed disparity condition, the positions of the focal region were swapped between the left and the right eyes the depth plane of the lateral motion was undefined (since the lines of sight would not meet at any point).

In different trials, the dots either jumped to the right in both eyes or to the left. To control for the final position of the target regions, the MID trials were also run with two variants – one in which the final position of the focal regions corresponded to their final position after the ‘left jump’ in the constant depth condition and one in which it corresponded to the final position of the ‘right jump’ in the constant depth condition. There were thus four different conditions which were presented to six mantises in interleaved trials: MID-left, MID-right, Constant-left and Constant-right. Each of these conditions were presented with the focal regions in each eye having either crossed or uncrossed disparity.

Each experimental run consisted of five interleaved replicates of every combination of four conditions and two disparity positions making for a total of forty trials. Two experimental runs were run for each of six mantises making for a total of 80 trials and ten replicates of each combination of conditions per animal. Trials were separated by a pause of 60 seconds.

### Experiment 2: Combined looming and stereoscopic cues to motion-in-depth

In this experiment, we presented the mantises with different combinations of looming and binocular cues. IOVD cues agreed with CD cues in every case, so this experiment did not attempt to separate their contribution. The basic stimulus consisted of a dark disc against a light background in the central region of the screen. In all conditions, the disc had a short spiral motion in the centre of the screen. The spiral motion was a modified version of the stimulus used in Experiment 1 and in previous studies where the stimulus reliably elicited strikes in the praying mantis (Nityananda et al., 2016b; Nityananda et al., 2018). In this version of the stimulus, the amplitude of the spiral was restricted to the centre of the screen rather than beginning at the periphery. The equations describing the motion were the same as given in Experiment 1 except for Eqns. 4 and 5. These were instead defined by two new equations

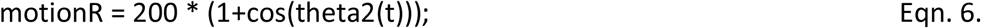

and

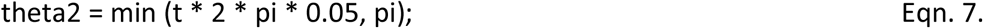

There were nine conditions in total (Fig. 3) which were presented in a randomised order in multiple experimental runs. Each experimental run consisted of 36 trials made up of four trials of each of the nine conditions. Each of ten mantises was presented with three experimental runs making for a total of 12 trials of every condition presented to each mantis. Trials were separated by a pause of 60 seconds to prevent habituation to stimulus presentation. The nine conditions were as follows:

1. CD-Loom: Here the stimulus had both changing disparity and size. The initial disparity and size (i.e. visual angle subtended) were set to simulate a target 20 cm away from the mantis. The stimulus subsequently increased in visual angle and changed disparity to simulate a target of 1 cm diameter at a distance of 2.5 cm in front of the mantis. The stimulus was simulated to move over 5 seconds with constant speed from 20 cm to 2.5 cm in front of the mantis. The change in visual angle and disparity were updated based on the simulated position of the target at any point in time. The stimulus thus had a diameter of 0.5 cm on the screen at the start and 4 cm at the end of the presentation. The visual angle subtended by the stimulus was 2.86° and 22.62° at the start and end of stimulus presentation respectively.
2. CD-SizeLarge: In this condition, the disparity changed as above. The visual angle was however kept constant to be the same as that subtended by a target of 1 cm diameter, 2.5 cm away from the mantis. The stimulus therefore had a diameter of 4 cm on the screen and subtended a visual angle of 22.62°.
3. CD-SizeSmall: This condition was the same as condition 2 above except that the visual angle by the stimulus disc was the same as a simulated target of 1 cm diameter, 10 cm away from the mantis. The stimulus therefore had a diameter of 1 cm on the screen and subtended a visual angle of 5.72°.
4. Loom-DispNear: In this condition, the target again loomed towards the mantis as described in condition 1. The disparity was, however, kept constant to simulate a target 2.5 cm away from the mantis.
5. SizeLarge-DispNear: Here both disparity and visual angle were kept constant to simulate a target 2.5 cm from the mantis. The stimulus size on the screen and the angle subtended were thus the same as in condition 2.
6. SizeSmall-DispNear: Both stimulus size and disparity were kept constant. The visual angle simulated a target 10 cm away from the mantis as described in condition 3. The disparity simulated a target 2.5 cm away from the mantis.
7. Loom-DispFar: In this condition, the target loomed towards the mantis as described in condition 1. The disparity was, however, kept constant to simulate a target 10 cm away from the mantis.
8. SizeLarge-DispFar: Both visual angle and disparity were kept constant. The visual angle simulated a target 2.5 cm away from the mantis while the disparity simulated a target 10 cm away from the mantis. The stimulus size on the screen and the angle subtended were thus the same as in condition 2.
9. SizeSmall-DispFar: Here both disparity and visual angle were kept constant to simulate a target 10 cm from the mantis. The stimulus size on the screen and the visual angle subtended were thus the same as in condition 3.

### Experiment 3: Motion-defined looming as a cue to depth

In a final experiment, we tested the contribution of internal motion to the perception of looming-based prey capture. The presence of a moving contrast edge has been shown to be critical to the perception of looming in insects. In this experiment, we asked if an edge defined by motion rather than contrast would also lead to strikes based on looming perception.

We used random-dot stimuli with the same background of dots as described for Experiment 2. We presented mantises with stimuli in two motion and two looming conditions (Fig. 5). In all conditions, the target moved with a spiralling motion in the centre of the screen as in Experiment 2 (Eqns. 1–3, 6 and 7). In the first motion condition, the target was defined by dots within the target region moving outward with a velocity of 2 pixels/s. As dots streamed outward, they were replaced from the centre to ensure the same density of dots was maintained. Once replaced at the centre, each dot was given a random direction along which it streamed outwards. Thus, in this condition, the target was defined by the outward internal motion forming a motion edge with the static background dots. In the second motion condition, there was no internal motion but the target consisted of a window within which a new set of dots was revealed behind the background layer of dots. This created the effect of a moving hole in the form of a drift-balanced stimulus.

Both of these motion conditions were presented as looming or not. In the looming condition, the size of the target simulated a target of diameter 1 cm looming from a distance of 20 cm away from the mantis to a distance of 2.5 cm away from the mantis as in the first condition of Experiment 2, i.e. increasing in angular size from 5.72° to 22.62°. In the non-looming condition, the size of the target was fixed and simulated a target of 1 cm diameter at a distance of 2.5 cm away from the mantis (22.62°). Experiments were run with either a crossed or uncrossed disparity. In the looming condition, crossed disparity cues across both eyes also changed to simulate a target approaching from 20 cm away to 2.5 cm away from the mantis. In the non-looming condition, crossed disparity cues simulated a target 2.5 cm from the mantis. In the uncrossed disparity conditions, the parallax between the focal regions was the same as in the crossed disparity conditions but the positions in the left and right eyes were swapped. Seven mantises were run in the crossed disparity condition and seven were run in the uncrossed disparity condition in separate experiments.

Mantises in each disparity condition were presented two motion conditions for each of two looming conditions, i.e., there were four stimulus conditions in each disparity condition. One experimental run consisted of an interleaved presentation of eight replicates of each of these four stimulus conditions with a total of 32 trials in an experimental run. Trials were separated by a pause of 60 seconds. Each mantis had two experimental runs making for a total of 72 trials and 16 replicates per combination of motion and looming conditions per mantis.

### Video and Statistical Analysis

All responses of the mantises were recorded using a Kinobo USB B3 HD Webcam (Point Set Digital Ltd, Edinburgh, Scotland) placed directly underneath the mantis. The camera was positioned so the screen could not be seen in the recording and all videos were blind to the stimulus condition. The parameters for every stimulus condition were saved separately during the experiment. The number of tracks, strikes and tensions made in each video were coded blind and this was saved separately. Tracks were sharp saccadic movements of the head in response to stimuli, strikes were rapid extensions of the forelegs and tensions involved a tensing of the forearms towards making strikes that were unreleased. The numbers of each of these were then matched to the parameters and the probability of response to each combination of parameters was calculated.

For all experiments we based our calculations of adequate power and the related minimum sample size on previous experiments (Nityananda et al., 2016b). Based on these previous results, we obtained an expected effect size (Cohen’s D) of 3.6. Such a high effect size implies that for a power of 0.8 in each experiment we would need a smaller minimum sample size of 5 animals and all experiments use a sample greater than this. Experiments used a within-subject experimental design and there was therefore no need for randomization between treatments. Experiment 3 had different animals in the crossed and uncrossed conditions and animals were assigned alternately to each of these treatments.

All analyses used generalized linear mixed models to analyse the data. The dependent variable was the probability of striking and the individual identity of the mantis was used as a random effect. Since the probability of striking was a binary decision (yes or no) we modelled the data using a binomial logistic link function. All data were analysed with the statistic software RStudio (version 1.1.383).

Experiment 1: Results from the MID-left and MID-right conditions were pooled as were results from the Constant-left and -right conditions. Motion condition (MID or Constant) was built into the model as a factor as was Disparity. Models were run with or without interaction between these two factors. Models were compared on the basis of the Bayesian Information Criterion (BIC).

Experiment 2: Size and Disparity were built in as factors into the model with each having three levels. The three levels for Size were called Loom, Large and Small, each respectively coding the conditions where the stimulus size increased as it loomed towards the mantis or was held constant to subtend an angle of 22.62° or 5.62°. The three levels for Disparity were called Changing Disparity, Near and Far each respectively coding the conditions where the stimulus disparity changed to simulate a target approaching the mantis or was held constant to simulate a target 2.5 cm or 10 cm away from the mantis. Models were run with or without interaction between these factors and animal identity was specified as a random effect. Models were compared on the basis of the Bayesian Information Criterion (BIC).

Experiment 3: Data were modelled with disparity, motion condition and looming condition as factors. Models were run with or without interaction between these factors. Models were compared on the basis of the Bayesian Information Criterion (BIC).

## Author Contributions

VN and JCAR designed the experiments. VN and GT programmed the stimuli. VN, CJ and JT ran the experiments. VN analysed the data. VN and JCAR wrote the paper.

## Acknowledgements

VN and GT were funded by Leverhulme Trust grant Research Leadership Award RL-2012–019 to JCAR. CJ carried out the work as part of a dissertation for the MSc from the Université de Rennes 1, France.

## Competing Interests

No competing interests declared.

## Funding

VN and GT were funded by Leverhulme Trust grant Research Leadership Award RL-2012–019 to JCAR.

## Data Availability

All data will be publically available on publication on figshare.com. Reviewers can access the data supporting this paper using the following private link: https://figshare.com/s/586b4f0e9b06deb8e253

